# Necrosis drives susceptibility to *Mycobacterium tuberculosis* in Polg^D257A^ mutator mice

**DOI:** 10.1101/2024.07.17.603991

**Authors:** CJ Mabry, CG Weindel, LW Stranahan, JJ VanPortfliet, JR Davis, EL Martinez, AP West, KL Patrick, RO Watson

**Affiliations:** Department of Microbial Pathogenesis and Immunology, Texas A&M Health, College of Medicine, Bryan, TX 77807, USA; Department of Veterinary Pathobiology, Texas A&M College of Veterinary Medicine and Biomedical Sciences, College Station, TX 77843, USA; Department of Medicine, Division of Infectious Diseases, Vanderbilt University Medical Center, Nashville, TN 37232; The Jackson Laboratory, Bar Harbor, Maine 04609, USA; Department of Pathology, Microbiology, and Immunology, Vanderbilt University Medical Center, Nashville, TN 37232

**Author notes:** Address Correspondence to Robert O. Watson.

## Abstract

The genetic and molecular determinants that underlie the heterogeneity of *Mycobacterium tuberculosis* (Mtb) infection outcomes in humans are poorly understood. Multiple lines of evidence demonstrate that mitochondrial dysfunction can exacerbate mycobacterial disease severity and mutations in some mitochondrial genes confer susceptibility to mycobacterial infection in humans. Here, we report that mutations in mitochondria DNA (mtDNA) polymerase gamma (POLG) potentiate susceptibility to Mtb infection in mice. Polg^D257A^ mutator mtDNA mice fail to mount a protective innate immune response at an early infection timepoint, evidenced by high bacterial burdens, reduced M1 macrophages, and excessive neutrophil infiltration in the lungs. Immunohistochemistry reveals signs of enhanced necrosis in the lungs of Mtb-infected Polg^D257A^ mice and Polg^D257A^ mutator macrophages are hyper-susceptible to extrinsic triggers of necroptosis *ex vivo*. By assigning a role for mtDNA mutations in driving necrosis during Mtb infection, this work further highlights the requirement for mitochondrial homeostasis in mounting balanced immune responses to Mtb.

## INTRODUCTION

Defects in mitochondria are associated with a variety of human diseases and metabolic disorders (1). Inherited mitochondrial diseases, caused by mutations harbored in the mitochondrial genome (mtDNA) or in nuclear-encoded mitochondrial genes, lead to devastating neurodegenerative and cardiovascular disorders (2). They are also frequently associated with chronic inflammation and autoimmunity. Type I interferonopathies, for example, are often caused by mutations that result in aberrant release of mitochondrial nucleic acids and chronic engagement of cytosolic nucleic acid sensing (3-5). Consistent with important links between mitochondrial health and immune function, patients with mitochondrial mutations are prone to frequent viral and bacterial infections that are often severe (6-7). They can also suffer from systemic inflammatory response syndrome (SIRS) caused by hyper-inflammation in the absence of infection (8). Furthermore, human SNPs in mitochondrial genes have been shown to confer susceptibility to several bacterial pathogens, including *Salmonella entrica* serovar Tyhi (9), *Listeria monocytogenes* (10), and *Mycobacterium leprae* (11). Despite clear connections between mitochondria and immunity, the mechanistic contributions of specific mitochondrial-associated mutations in driving susceptibility to infection remain unclear.

*Mycobacterium tuberculosis* (Mtb) is the causative agent of tuberculosis, an ancient disease that remains the number one infectious disease killer worldwide (12). The success of Mtb is driven in large part by its ability to manipulate and establish a replicative niche in host innate immune cells, primarily macrophages (13). A key hub in the Mtb-macrophage host-pathogen interface are mitochondria, which maintain cellular energetics, control lipid metabolism, and generate reactive oxygen species, as well as gatekeep entry into programmed cell death (14). While the precise molecular determinants of TB disease severity remain elusive, decades of research have identified a role for cell death in dictating TB disease outcomes. Apoptosis, or “immunologically silent” cell death correlates with protection, whereas necrotic forms of cell death are associated with mycobacterial cell-to-cell spread and immunopathology (15-18). Consequently, TB has evolved a variety of strategies to inhibit apoptosis while promoting necrosis (19).

Several studies have mechanistically shown how distinct modes of cell death (e.g. inflammasome-mediated pyroptosis, RIPK-dependent necroptosis, ferroptosis) promote Mtb pathogenesis (20-24). We and others have begun to link cell death during Mtb infection with disruptions in mitochondrial homeostasis (14, 25-26). We recently reported that mutations in the mitochondrial-associated protein leucine-rich repeat kinase 2 (LRRK2) can promote necroptosis during Mtb infection by rendering mitochondrial membranes susceptible to damage by the pore-forming protein gasdermin D, resulting in the release of mitochondrial ROS (26). Building on this finding, we set out to determine how other mitochondrial mutations alter Mtb infection outcomes.

Here, we investigate how Mtb infection proceeds in a common model of mitochondrial dysfunction, the Polg mutator mouse. These mice contain a knock-in D257A mutation in the exonuclease domain of Polg, the sole mitochondrial DNA polymerase (27). This mutation disrupts the proofreading ability of Polg and subsequently results in accumulation of mtDNA mutations in these mice as they age (27). Although this mouse model does not harbor a specific human disease-associated mutation, it presents with symptoms that parallel many of the clinical manifestations seen in mitochondrial disease patients, such as cardiomyopathy, progressive hearing loss, and anemia (28). These mice also have exacerbated inflammatory responses and are susceptible to septic shock, which correlates with clinical symptoms seen in mitochondrial disease patients (3). They have been a useful tool for studying mitochondrial dysfunction in aging (27, 29), but only a few studies have used them to study infection (30). Here, we report that Polg^D257A^ mutator mice display enhanced susceptibility to Mtb infection evidenced by higher bacterial burdens, excessive neutrophil infiltration in the lung, and enhanced necrosis. These findings argue that mtDNA fidelity is required to maintain balanced cell death outcomes during Mtb infection.

## RESULTS

### Polg^D257A^ mutator mice harbor higher Mtb bacterial burdens

To better understand how mitochondrial-associated gene mutations drive Mtb infection outcomes, we chose to study the Polg^D257A^ mutator mouse model (B6.129S7(Cg)-Polg^tm1Prol^/J). These mice accumulate linear mtDNAs and fragmented mtDNA genomes that contribute to mitochondrial dysfunction and aging (27). Using a low-dose aerosol model, which administers 100-200 Mtb bacilli into the lungs of each animal, we infected WT (n=6) and Polg^D257A^ mutator (n=9) mice at approximately 3 months of age. We chose to infect young mice to avoid variability in mtDNA instability and the potential for inflammation-associated pathology in our cohort (Polg^D257A^ mutator mice experience premature aging and early death (27-29)). At 21 days post-infection, we sacrificed mice, homogenized a portion of the lungs, spleen, and liver and enumerated Mtb colony-forming units (CFUs) to measure bacterial burden. We observed a significant increase (half-log) in Mtb recovered from the lungs of Polg^D257A^ mutator mice compared to WT mice (**Figure 1A**). Polg^D257A^ mutator mice were also more permissive to Mtb dissemination, as bacterial burdens were higher in their spleens and livers (**Figure 1B-C**). These findings demonstrate that mice experiencing Polg-dependent mitochondrial dysfunction cannot properly control Mtb replication and spread *in vivo*.

**Figure 1.**
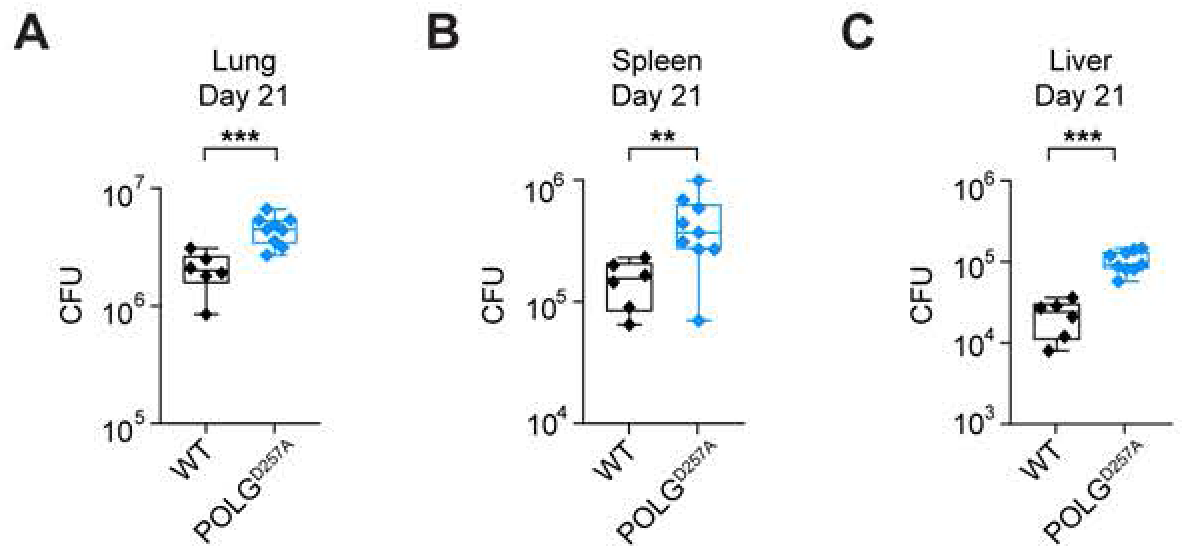
Polg^D257A^ mutator mice display increased Mtb replication and dissemination *in vivo*. Mtb colony forming units (CFUs) collected from WT and Polg^D257A^ mice lungs **(A)**, spleens **(B)**, and livers **(C)** at day 21 post-Mtb infection. Statistical analysis: *p < 0.05, **p < 0.01, ***p < 0.001, ****p<0.0001. Statistical significance determined for (A-C) using Mann-Whitney U test.

### Polg^D257A^ mutator mice lungs exhibit increased neutrophils and fewer M1 macrophages during Mtb infection

To grossly examine the inflammatory state of the lungs in Mtb-infected Polg^D257A^ mutator mice, we evaluated lung tissues via hematoxylin and eosin (H&E) staining. As is characteristic of Mtb infection, we observed histiocytic inflammation (aggregation of monocyte- and macrophage-derived cells) in the lungs of both genotypes (**Figure 2A-B).** Curiously, despite harboring higher bacterial loads, Polg^D257A^ mutator mice displayed less histiocytic lung inflammation at day 21 post-infection compared to WT (**Figure 2C,** black outline in H&E images). This suggests that Polg^D257A^ mutator mice fail to effectively recruit one or more types of immune cells into infected regions of the lung. In contrast, we found that Polg^D257A^ mutator mice lungs were more prone to neutrophil cluster formation, a phenomenon typically associated with NET release and pathogen clearance (31,32) (**Figure 2D**, white outline in H&E images). These findings argue that an altered immune cell landscape dominates the lungs of Polg^D257A^ mutator mice during Mtb infection.

**Figure 2.**
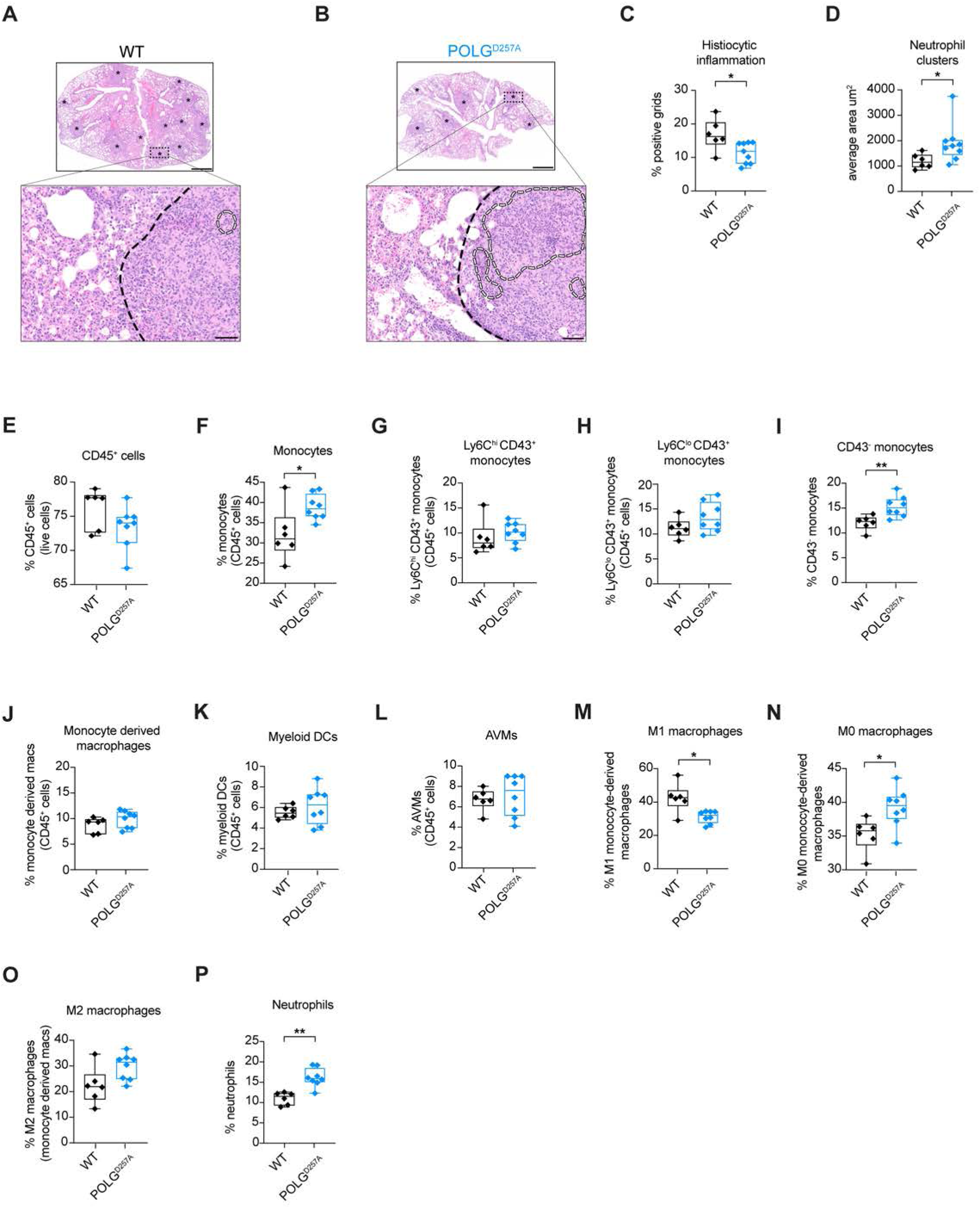
Polg^D257A^ mutator mice display increased neutrophils and less M1 macrophages during Mtb infection. Hematoxylin and eosin staining of WT **(A)** and Polg^D257A^ **(B)** mice lungs at day 21 post-Mtb infection. Asterisks denote areas of consolidated and infiltrated by inflammatory cells. Black dashed line delineates regions of healthy versus inflamed areas of lung in right panel. Outlined regions (white dashed lines) in denote areas of neutrophil clusters. Upper panel scale bar is 800 μM. Lower panel scale bar is 20 μM. **(C)** Quantification of histiocytic inflammation using H&E staining of WT and Polg^D257A^ mice lungs at day 21 post-Mtb infection. Each data point represents the percent of positive grids with histiocytic inflammation in WT and Polg^D257A^ mice Mtb-infected lungs. **(D)** Average total surface area of neutrophil clusters using H&E staining of WT and Polg^D257A^ mice lungs at day 21 post-Mtb infection. For flow cytometry analysis, WT (n=6) and Polg^D257A^ (n=8) mice lungs were analyzed. Lung CD45^+^ cells **(E)**, monocytes **(F)**, Ly6C^hi^ CD43^+^ monocytes **(G)**, Ly6C^lo^ CD43^+^ monocytes **(H)**, CD43^-^ monocytes **(I)**, monocyte derived macrophages **(J)**, myeloid dendritic cells **(K)**, alveolar macrophages **(L),** M1 macrophages **(M)**, M0 macrophages **(N)**, M2 macrophages **(O)**, and neutrophils **(P)** in WT and Polg^D257A^ mice analyzed by flow cytometry at day 21 post-Mtb infection. Statistical analysis: *p < 0.05, **p < 0.01, ***p < 0.001, ****p <0.0001. Statistical significance determined for (C-P) using Mann-Whitney U test.

To further define innate immune cell populations in the lungs of Polg^D257A^ mutator mice during Mtb infection, we isolated single cell suspensions and used flow cytometry to identify key cell types. Overall, we measured similar numbers of hematopoietic cells (CD45^+^) in the lungs of Mtb-infected Polg^D257A^ mutator and WT mice (**Figure 2E**). However, the percentage of total monocytes, in particular classical Ly6C^+^ CD43^-^ monocytes, was higher in Polg^D257A^ mutator mice (**Figure 2F-I**). Previous studies have correlated CD43 with control of Mtb in macrophages, and in mice, the absence of CD43 leads to increased bacterial load and defective granuloma formation (33). We did not observe major differences in the percentage of monocyte-derived macrophages (CD45^+^ CD11b^+^ Ly6C^-^ SSC^hi^) (**Figure 2J**), dendritic cells (CD45^+^ CD11b^+^ CD11c^+^) (**Figure 2K**), or alveolar macrophages (CD45^+^ CD11b^low/neg^ CD11c^+^ CD170^+^) (**Figure 2L**).

Consistent with the lower levels of histiocytic inflammation (**Figure 2C**), we measured fewer M1 macrophages (CD86^+^ macrophages) in Mtb-infected Polg^D257A^ mutator mice (**Figure 2M**). This was a concomitant with increased percentage of M0 (CD86^-^ CD206^-^) macrophages (**Figure 2N**), suggesting a macrophage polarization defect in Polg^D257A^ mutator mice. M2 (CD86^-^ CD206^+^) macrophages were similar between the two genotypes (**Figure 2O**). Importantly, pro-inflammatory M1 macrophages play an important role in the early control of Mtb by promoting granuloma formation, triggering T cell infiltration, and upregulating antimicrobial molecules like nitric oxide (34-36). Concomitant with fewer M1 macrophages, we observed a significant increase in the recruitment of neutrophils (CD45*^+^* C220^-^ CD11b^+^ Ly6G^+^) in Polg^D257A^ mice lungs (**Figure 2P**). Enhanced neutrophil recruitment has been previously shown in animal models and tuberculosis patients to correlate with tissue damage and poor Mtb infection outcomes (37-39). Collectively, these results suggest that a combination of defective M1 macrophage polarization, low expression of CD43 on monocytes, and/or enhanced neutrophil infiltration result in higher Mtb bacterial burdens in Polg^D257A^ mice.

### Decreased M1 macrophage responses drive susceptibility to Mtb infection in Polg^D257A^ mutator mice

Previous studies have linked Polg-related mitochondria dysfunction to elevated type I IFN expression following exposure to pattern recognition receptor (PRR) agonists such as LPS and dsDNA (3). To begin to understand if such a defect contributes to Mtb susceptibility in Polg^D257A^ mutator mice, we isolated primary bone marrow derived macrophages (BMDMs) from WT and Polg^D257A^ mice and infected them with Mtb (MOI=5). We did not observe differences in the ability of Polg^D257A^ BMDMs to elicit *Ifnb1* or interferon-stimulated gene (ISG) transcript expression in response to Mtb (**Figure S2A**), nor did we measure differences in expression of Viperin (encoded by *Rsad2*) at the protein level 24h post-Mtb infection (**Figure S2B**). These data suggest that the cell-intrinsic ability of Polg^D257A^ macrophages to generate type I IFN in response to Mtb is intact *ex vivo*.

We next asked how cytokine expression was altered in Mtb-infected Polg^D257A^ mutator mice. In lung homogenates, ISGs *Il1ra*, *Isg15*, and *Rsad2* (which encodes Viperin) were expressed similarly in WT and Polg^D257A^ Mtb-infected lungs (**Figure 3A-C**). However, consistent with low M1 macrophage numbers (**Figure 2M**), pro-inflammatory transcripts like *Tnfa*, *Nos2*, *Ifng*, *Nlrp3*, *Cxcl9*, and *Cxcl10* were less abundant in the Polg^D257A^ lungs (**Figure 3D-I**). Other pro-inflammatory cytokines (*Il1b* and *Il6*) and chemokines (*Cxcl1 and Cxcl5*) were consistently expressed by both genotypes (**Figure S2C-F**). Because transcript levels do not always correlate with protein levels, we also performed western blot analysis on lung homogenates. Although most secreted cytokines are undetectable in lung homogenates, we were able to confirm lower expression of Nlrp3 at the protein level (**Figure 3J**). Curiously, although *Rsad2* and *Il1b* transcript abundance was similar in WT and Polg^D257A^ lungs, protein levels were, on average, lower in Polg^D257A^ (**Figure 3J**). Collectively, these data demonstrate that Polg^D257A^ mice generate a weak pro-inflammatory milieu in the lung during early Mtb infection.

**Figure 3.**
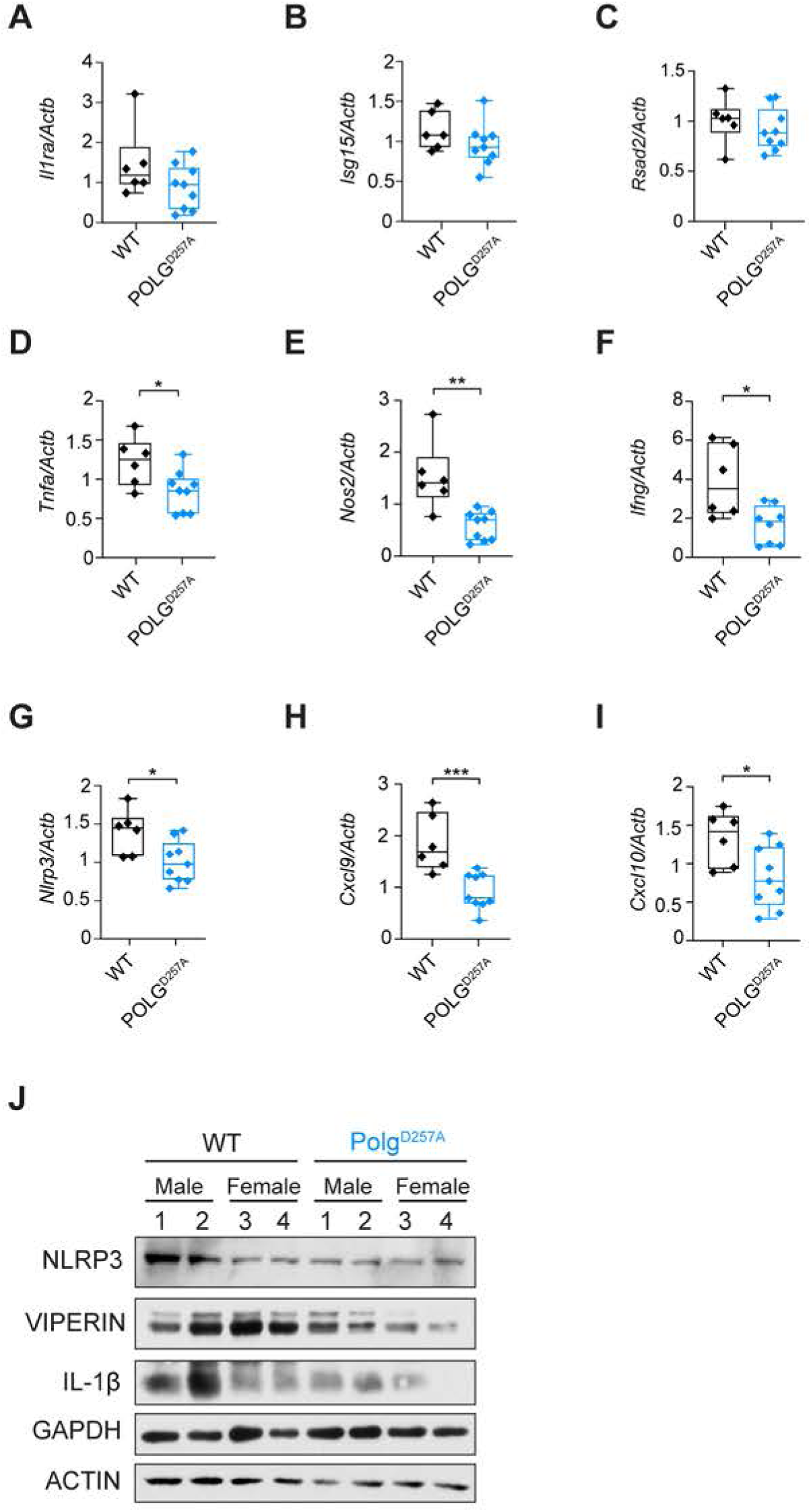
Dampened M1 macrophage pro-inflammatory response is a hallmark of Polg^D257A^ mutator mice during Mtb infection. Lung *Il1ra* **(A)**, *Isg15* **(B)**, *Rsad2* **(C)**, *Tnfa* **(D)**, *Nos2* **(E)**, *Ifng* **(F)** *Nlrp3* **(G)**, *Cxcl9* **(H),** *Cxcl10* **(I)** transcripts in WT and Polg^D257A^ mice analyzed by qRT-PCR at day 21 post-Mtb infection. Transcripts were normalized to *Actb* transcript levels. **(J)** Immunoblot analysis of NLRP3, VIPERIN, and IL-1β protein levels in WT and Polg^D257A^ mouse lungs at day 21 post-Mtb infection. GAPDH and ACTIN were used as a loading control. Statistical analysis: *p < 0.05, **p < 0.01, ***p < 0.001, ****p < 0.0001. Statistical significance determined for (A-I) using Mann-Whitney U test.

Lastly, looking beyond the lung, we measured circulating cytokines in Mtb-infected WT and Polg^D257A^ mice. Briefly, we used the ProcartaPlex™ Multiplex Immunoassay Kit to analyze serum isolated from WT (n=6) and Polg^D257A^ (n=9) mice. Most of the analytes in our 12-plex panel were below the level of detection. For the analytes we could measure (IFN-γ, TNF-α, CXCL5, CXCL9, and CXCL10), we observed no statistically significant differences between WT and Polg^D257A^ mice (**Figure S2G**). These data argue that at Day 21 post-Mtb infection, the Polg^D257A^ mutation drives immune changes in the lung without making a significant impact on the circulating immune milieu.

### Necrotic debris dominates Polg^D257A^ mutator mouse lungs during Mtb infection

Having observed altered immune cell infiltration and inflammatory transcript abundance in Mtb-infected Polg^D257A^ mutator mice, we next looked more carefully at immunopathology in the lungs. We and others have previously associated necrotic debris, characterized by neutrophil clusters with granular nuclear debris from dead cells, with poor tuberculosis outcomes in mice (26, 40). Accordingly, we observed increased clusters of necrotic debris in Mtb-infected Polg^D257A^ mutator mice lungs compared to WT (**Figure 4A-C, white arrows**). Because necrosis is associated with exuberant Mtb growth and virulence (22, 40), we next used acid-fast staining to look for Mtb bacilli in areas of consolidated inflammatory infiltrates **(Figure 4D-E, black outline)**. We denoted instances of growth characteristic of cording, a virulence-related form of Mtb growth associated with necrosis, specifically in the lungs of Polg^D257A^ mutator mice (**Figure 4D-E**) (19, 40). To quantify the number of Mtb bacilli in areas of neutrophil clusters and necrotic debris, we developed an unbiased image analysis workflow that scored the number of AFB-positive bacilli per selected fields of view. Using this approach, we measured significantly more Mtb bacilli in necrotic regions of Polg^D257A^ mutator mice lungs compared to WT (**Figure 4F**). These findings suggest that areas of necrotic debris that dominate Polg^D257A^ mutator mice lungs correlate with a failure to restrict Mtb growth.

**Figure 4.**
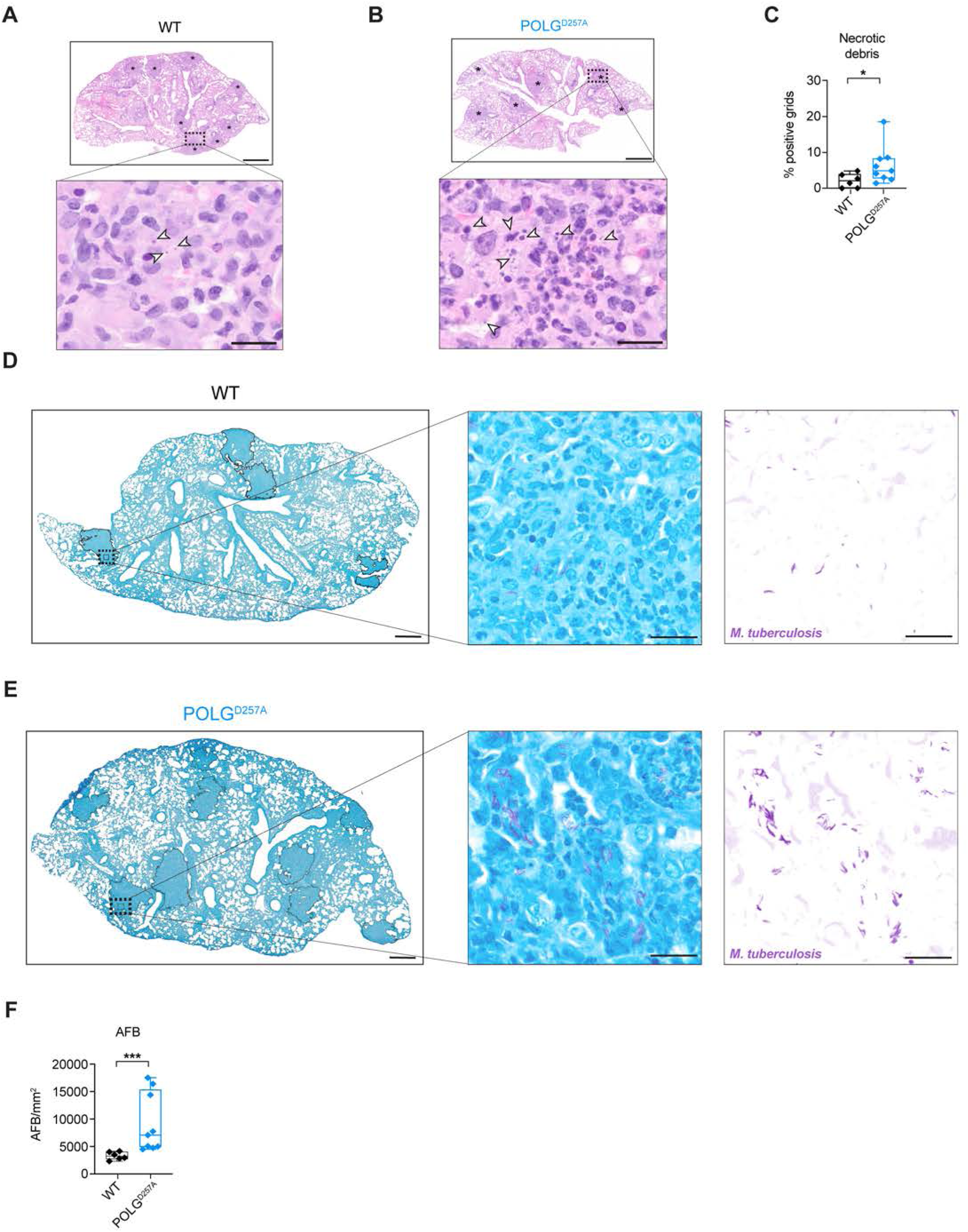
Polg^D257A^ mutator mice exhibit increased necrotic debris and higher bacterial burden in areas of necrosis during Mtb infection. Hematoxylin and eosin staining of WT **(A)** and Polg^D257A^ **(B)** mice lungs at day 21 post-Mtb infection. Asterisks denote areas of consolidated and infiltrated by inflammatory cells. Outlined regions (white dashed lines) denote areas of necrotic debris. White arrows denote necrotic debris (extracellular, circular, densely structures of dead cells). Upper panel scale bar is 800 μM. Lower panel scale bar is 10 μM. **(C)** Quantification of necrotic debris using H&E staining of WT and Polg^D257A^ mice lungs at day 21 post-Mtb infection. Each data point represents the percent of positive grids with necrotic debris in WT and Polg^D257A^ mice Mtb-infected lungs. Acid-fast bacilli (AFB) staining of WT **(D)** and Polg^D257A^ **(E)** mice lungs at day 21 post-Mtb infection. Black outlines denote consolidate areas of inflammatory infiltrates in left panel. The middle and right panels represent insets of AFB staining and color deconvolution. Left panel scale bar is 800 μM. Middle and right panels scale bar is 20 μM. **(F)** Quantification of AFB bacilli in black outlined consolidate areas of inflammatory infiltrates (See methods section titled “Quantification of acid-fast bacteria (AFB) staining”) Statistical analysis: *p < 0.05, **p < 0.01, ***p < 0.001, ****p < 0.0001. Statistical significance determined for (C,F) using Mann-Whitney U test.

### Polg^D257A^ macrophages are prone to necrotic death and cell death in response to Mtb infection

To better understand the molecular mechanisms underlying the susceptibility of Polg^D257A^ mutator mice, we asked if their propensity to accumulate necrotic debris was observable only in the infected lung, or if there was a cell-intrinsic propensity for Polg^D257A^ macrophages to undergo necrosis. To test this, we isolated bone marrow and differentiated macrophages from WT and Polg^D257A^ mutator mice at either 6 or 12 months of age. These ages represent the typical “onset” and “late” stages of mitochondrial disease in Polg^D257A^ mutator mice as they relate to human patients (27, 41). We first infected WT and Polg^D257A^ BMDMs from 6-month-old mice with Mtb (MOI=5) and tracked cell death kinetics using propidium iodide (PI). After 24h of infection, Polg^D257A^ macrophages were more prone to Mtb-induced cell death than WT controls (**Figure 5A**). Because Mtb infection triggers several cell death pathways in macrophages (apoptosis (42), ferroptosis (24), and inflammasome-mediated pyroptosis (20-21)) and PI cannot distinguish between these forms of cell death, we next exposed cells to canonical cell death triggers to gain additional insight into the type of cell death that is enhanced in Polg^D257A^ BMDMs. To begin, we treated WT and Polg^D257A^ BMDMs from 6-month-old mice with the apoptosis inducer staurosporine, a protein kinase inhibitor that triggers apoptosis via aberrant caspase activation. In response to a low dose of staurosporine (100 nm), we observed slightly more propidium iodide incorporation in Polg^D257A^ BMDMs (**Figure 5B**). These findings are consistent with previous work that found increased apoptotic markers associated with accelerated aging in Polg^D257A^ mutator mice (24). Next, to induce necroptosis, we treated WT and Polg^D257A^ BMDMs with recombinant murine TNFα in combination with a SMAC mimetic and pan-caspase inhibitor to initiate RIPK1/RIPK3/MLKL-dependent necroptosis (43). Polg^D257A^ macrophages isolated from 6-month-old mice were highly susceptible to necroptosis (**Figure 5C**) and this phenotype was enhanced in Polg^D257A^ BMDMs from 12-month-old mice (**Figure 5D**). These findings suggest that Polg^D257A^ macrophages are more sensitized to necrotic forms of cell death, which may account for enhanced necrosis and Mtb dissemination *in vivo*.

**Figure 5.**
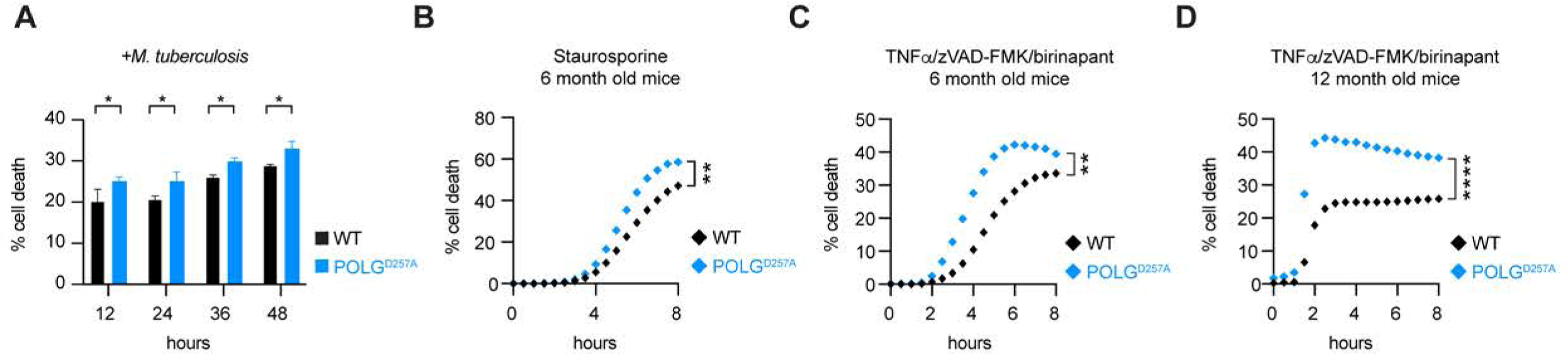
Polg^D257A^ mutator macrophages are prone to necrotic cell death. **(A**) Cell death in 6-month-old WT and Polg^D257A^ macrophages infected with Mtb for 48h (MOI=5). **(B)** Cell death in 6-month-old WT and Polg^D257A^ macrophages over a time course of apoptosis induction with 100nm staurosporine. Cell death in 6-month-old **(C)** or 12-month-old **(D)** WT and Polg^D257A^ macrophages treated with 100 ng/ml TNFα, 500 nM Birinapant, and 20 μM zVAD-FMK to induce necrosis. Cell death was measured by propidium iodide uptake (% cell death= Propidium iodide positive/ total cell count *100%). Statistical analysis: n=3 *p < 0.05, **p < 0.01, ***p < 0.001, ****p<0.0001. Statistical significance determined for (A) using one-way ANOVA with Sidak’s post-test and (B-D) using two-way ANOVA with Tukey’s post-test.

## Discussion

Because mitochondrial-associated mutations can increase the risk of life-threatening infections, there is a critical need to understand how altered mitochondrial homeostasis affects the immune response to pathogens. Here, we describe how a mutation in mitochondrial POLG, which results in the accumulation of mtDNA mutations and mtDNA instability, confers susceptibility to Mtb infection by enhancing necrosis. We demonstrate that in response to Mtb, Polg^D257A^ mutator mice fail to mount protective immune responses in the lung. We propose this stems in large part from the propensity of Polg^D257A^ macrophages to preferentially undergo necrosis over other forms of programmed cell death when infected with Mtb. Enhanced necrosis is a likely driver for the pro-bacterial cellular immune milieu (high neutrophils, low M1 macrophages) we observe in Polg^D257A^ mutator mice. It may also directly support high levels of Mtb growth in Polg^D257A^ lungs. Studies from the Ramakrishnan lab have repeatedly linked necrosis to extracellular mycobacterial cording and poor disease outcomes in the zebrafish model (44-45). In human macrophages, Mtb replication is enhanced in cells undergoing late stages of necrosis that exhibit evidence of plasma membrane damage (46). Mtb has also been shown to thrive in macrophages that have phagocytosed Mtb-infected PMNs that died via necrosis (40). Although we are unable to decisively assess whether Mtb bacilli in Polg^D257A^ lungs are extracellular due to limitations of H&E and AFB staining, our data convincingly correlate enhanced necrosis in Polg^D257A^ mutator mice with high Mtb bacterial burdens.

Over the course of these studies, we were surprised to find that despite harboring more bacteria in the lungs, spleen, and liver (**Figure 1**), Polg^D257A^ mutator mice expressed lower levels of cytokines/chemokines overall (**Figure 3 and S2**). It is possible that the accumulation of mtDNA mutations and chronic stimulation of pattern recognition receptors by mitochondrial DAMPs render Polg^D257A^ immune cells refractory to immune signaling upon pathogen sensing (47). We previously reported a phenotype like this in LRRK2 KO macrophages, which have high basal type I IFN expression and are unable to induce ISG expression in response Mtb (48). It is also possible that the failure of Polg^D257A^ mutator mice to upregulate pro-inflammatory cytokines stems from rewiring of cellular metabolism and mitochondrial OXPHOS, which is uniquely regulated in Mtb-infected macrophages (49-50). Previous work has linked the Polg^D257A^ mutation to altered type I IFN signaling in response to bacterial pathogens. Specifically, the West lab reported that 6-48h post-intratracheal instillation of *Pseudomonas aeruginosa* strain O1 (PAO1), Polg^D257A^ mutator mice exhibit elevated levels of ISGs in the lung and BAL fluid (30). It is possible that upon initial encounter with Mtb, Polg^D257A^ mutator mice similarly hyperinduce ISGs and pro-inflammatory cytokines and that we are reporting on a different, chronic response at 21 days post-infection. However, it is also possible that the Polg^D257A^ mutation responds to and elicits an immune milieu to different pathogens in different ways. Indeed, we know that *Pseudomonas* and Mtb initially elicit type I IFN expression through distinct signaling pathways (TLR4/TRIF (51) and cGAS (52), respectively) and likely interface with mitochondria in distinct ways as well. Investigations of Mtb-infected Polg^D257A^ mutator mice at additional early time points may help us distinguish between these two potential explanations.

These studies were motivated, in part, by several genome wide association studies (GWAS) that identified SNPs in mitochondrial genes that confer susceptibility to mycobacterial infection in humans. Most of these studies looked at patients with leprosy, which is caused by *Mycobacterium leprae*, a close cousin of Mtb. Mutations in genes like *Park2*, which encodes the mitophagy factor Parkin (53-54), *Tfam*, which encodes a histone-like protein of the mitochondrial genome (11), as well as *Polg* (11) have been linked to leprosy susceptibility or severity of disease. The Mtb phenotype we report is consistent with GWAS that identified a human SNP in POLG associated with multibacillary leprosy—a form of the disease characterized by high bacterial loads in lesions (11). Collectively, our study bolsters confidence in these GWAS and provides new insight into the potential mitochondria-associated mechanisms driving altered susceptibility to mycobacterial disease.

### Limitations of the study

One drawback of the Polg^D257A^ mutator mouse model is that the mice accumulate mutations throughout the entire mtDNA genome over time (55). Thus, one cannot ascribe pathological phenotypes to a specific defect in a single mitochondrial gene. Likewise, although human patients with mutations in the exonuclease “proofreading” domain of POLG have been linked to diseases such as Alpers and Leigh syndrome (56, 57), Polg^D257A^ mutator mice do not mirror a specific model of mitochondria disease. Given these caveats, future investigations may wish to probe Mtb pathogenesis in mice that more accurately model mitochondrial mutations in humans. Likewise, it will be interesting to determine the susceptibility of the Polg^D257A^ mutator mice at later timepoints during Mtb infection since mtDNA mutations accumulate with age in these mice. Our bacilli staining in particular hints at a bifurcation in our mouse data, whereby certain mice harbor incredibly high Mtb burdens in necrotic lesions (**Figure 4**). It is tempting to speculate that these are mice that have accumulated the most mtDNA mutations—a hypothesis that can be tested by sequencing mtDNA from the lungs of the Polg^D257A^ mutator mice. Despite these limitations, our study adds to a growing literature arguing that mitochondrial health is required to generate protective immune responses to Mtb. Accordingly, we propose that disruption to mitochondrial homeostasis, via mutation, metabolic disease, stress, etc. plays a critical role in dictating the spectrum of tuberculosis disease in humans.

## Acknowledgements

We would like to thank the members of the Watson and Patrick labs for their critical review and feedback in the preparation of this manuscript. We thank the UTHSC Regional Biocontainment Laboratory (NIH Research Grant number UC6 AI058616) staff, Dr. Kellie Brown, for their excellent support for MagPix assays. UTHSC facility equipment was supported in part by NIH Research Grant numbers UC7AI180313 and G20AI167349. We thank Dr. Samantha Bell, Assistant Professor of Microbiology, Biochemistry & Molecular Genetics at Rutgers New Jersey Medical School, for her contributions to this story at its early stages. Funding was provided by NIH/NIAID to R.O.W. and K.L.P. (R01AI155621), W81XWH-20-1-0150 to A.P.W. from the Office of the Assistant Secretary of Defense and for Health Affairs through the Peer Reviewed Medical Research Programs, NIH/NIAID to C.J.M. (F31AI176795), NIH training grant T32GM135748 to E.L.M., NIH/NIAID to E.L.M. (F31AI176821), NIH/NRSA to J.J.V. (F31AI179168), and the Parkinson’s Foundation Launch Award PF-Launch-938138 (C.G.W.).

## Author contributions

Conceptualization, C.J.M., K.L.P., and R.O.W.; Investigation, C.J.M., C.G.W., L.W.S., J.J.V., J.R.D., E.L.M., A.R.W., K.L.P, and R.O.W.; Methodology, C.J.M, C.G.W., L.W.S., J.J.V., J.R.D., E.L.M., A.R.W., K.L.P., and R.O.W.; Visualization, C.J.M., K.L.P, and R.O.W.; Writing, C.J.M., K.L.P., and R.O.W.; Supervision, R.O.W. and K.L.P.

## MATERIALS and METHODS

### Mouse husbandry and strains

*POLG^D257A/D257A^* C57BL/6J mutator mice were purchased from the Jackson laboratory. *POLG^D257A/+^* breeder pairs used to generate *POLG^+/+^*and *POLG^D257A/D257A^* experimental mice were obtained from male *POLG^D257A/+^* to female C57BL/6J crosses. All mice used in experiments were compared to age- and sex- matched controls and fed 4% standard chow. Littermate controls were used in all experiments. For *ex vivo* BMDM experiments, male mice 6-month and 12-month of age were used. For *in vivo* Mtb infections, mice were infected at 12 weeks of age. All animals were housed, bred, and studied at Texas A&M Health Science Center under approved Institutional Care and Use Committee guidelines.

### M. tuberculosis

The Erdman strain was used for all *M. tuberculosis* infections. Low passage lab stocks were thawed for each experiment to ensure virulence was preserved. *M. tuberculosis* was cultured in roller bottles at 37°C in Middlebrook 7H9 broth (BD Biosciences) supplemented with 10% OADC (BD Biosciences), 0.5% glycerol (Fisher), and 0.1% Tween-80 (Fisher) or on 7H10 plates. All work with *M. tuberculosis* was performed under Biosafety level 3 containment using procedures approved by the Texas A&M University Institutional Biosafety Committee.

### Primary cell culture

Bone marrow derived macrophages (BMDMs) were isolated by flushing mouse femurs and tibias with 10 mL DMEM 1 mM sodium pyruvate. Cell suspensions were centrifuged at 400 rcf for 5 min and responded in BMDM media (DMEM, 20% FBS (MilliporeSigma), 1 mM sodium pyruvate (Lonza), 10% MCSF conditioned media (Watson lab)). Cells were plated on non-TC treated plates and differentiated at 37°C 5% CO_2_. On day 3 following plating, cells were fed 50% additional volume with BMDM media. Cells were harvested at day 7 with 1XPBS EDTA (Lonza).

## METHOD DETAILS

### M. tuberculosis infection

For *M. tuberculosis* infections *ex vivo*, low passage Mtb was prepared by growing it to log phase (OD600 0.6-0.8). Bacterial cultures were spun at 58 rcf for 5 min to remove large clumps. The bacteria was then pelleted at 2103 rcf for 5 min and washed with 1XPBS. The wash step was repeated twice. The resuspended bacteria cultures were sonicated at 70% amplitude for 10 sec repeated three times (Branson Ultrasonics Corp.) followed by a low speed spin (58 rcf) to remove remaining clumps. The bacteria were diluted in DMEM (Hyclone) + 10% horse serum (Gibco) for *ex vivo* infections, or 1X PBS for *in vivo* infections. For *ex vivo* infections, plates containing cells and bacteria were spun at 234 rcf for 10 min to synchronize infection. Fresh BMDM media was then added to the cells. See “plate based assays for cell death” for details on measuring cell death kinetics during Mtb infection.

### Plate based assays for cell death

BMDMs were plated in 96-well clear bottom plates (Corning) at 2.5×10^4^ cells/well in 50 μL of media. Following cell adherence (1 hr), an additional 25 μL was added to each well. The following day, media was removed, and respective agonists were used to initiate various cell death modalities. For cell death assays, 5 μg/ml propidium iodide (PI) (ThermoFisher) was added at the time of adding cell death agonists. Total cell numbers used for normalization were counted on a subset of cells using NucBlue (ThermoFisher) in 1X PBS. Live cell imaging was done using 4X magnification on either a Lionheart plate reader or Cytation 5 (Biotek). Post-run analysis was conducted using Gen5 v3.15 (Biotek) software. For necroptosis induction, cells were activated with 100 ng/ml recombinant murine TNFα (PeproTech), 500 nM SMAC mimetic birinapant (Selleck Chemicals), and 20 μM pan-caspase inhibitor z-VAD-FMK (R&D Systems). Apoptosis was induced with 100 nM stauroporine (Tocris Bioscience).

### *In vivo* Mtb infections

All infections were performed using procedures approved by Texas A&M University Institutional Care and Use Committee. The Mtb inoculum was prepared as described above. Age- and sex-matched mice were infected via inhalation exposure using a Madison chamber (Glas-Col) calibrated to introduce 100-200 CFUs per mouse. For each infection, approximately 3 mice were euthanized immediately, and their lungs were homogenized and plated to verify an accurate inoculum. Infected mice were housed under BSL3 containment and monitored daily by lab members and veterinary staff. At the indicated time points, mice were euthanized, and tissue samples were collected. Blood was collected in serum collection tubes, allowed to clot for 1-2 hrs at RT, and spun at 1000 rpm for 10 min to separate serum. Organs were divided to maximize infection readouts CFUs: left lobe lung and 1⁄2 spleen; histology: 2 right lung lobes and 1⁄4 spleen; RNA: post-caval lobe and 1⁄4 spleen; Flow: right inferior lobe). For histological analysis organs were fixed for 24 hrs in 10% neutral buffered formalin and moved to ethanol (lung, spleen). Organs were further processed as described below. For cytokine transcript analysis, organs were homogenized in Trizol, and RNA was isolated as described below. For CFU enumeration, organs were homogenized in 5 ml PBS + 0.1% Tween-80, and serial dilutions were plated on 7H10 plates. Colonies were counted after plates were incubated at 37°C for 3 weeks.

### Multiplex immunoassay

The levels of a panel of cytokines and chemokines in serum were measured using a 12-plex ProcartaPlex™ Multiplex Immunoassay according to manufacturer’s instructions (Invitrogen, Carlsbad CA, USA). Standards were prepared to determine the concentration of cytokines and chemokines in the samples. The samples were run on a Magpix instrument and analyzed with Milliplex Analyst v5.1 software. For data analysis, a five-parameter logistic curve fitting method was applied to the standards and the sample concentrations extrapolated from the standard curve. All experiments using infectious agents were conducted in BSL-3 or ABSL-3 facilities of the University of Tennessee Health Science Center Regional Biocontainment Laboratory (UTHSC RBL) compliant with protocols that were reviewed and approved by the UTHSC Institutional Biosafety Committee (IBC) and the Institutional Animal Care and Use Committee (IACUC).

### Gene expression analysis by RT-qPCR

For lung tissue, RNA was isolated using Direct-zol RNAeasy kits (Zymogen). cDNA was synthesized with Bio-Rad iScript Direct Synthesis kits (BioRad) per manufacturer’s protocol. RT-qPCR was performed in triplicate wells using PowerUp SYBR Green Master Mix. Data was analyzed on a QuantStudio 6 Real-Time PCR System (Applied Biosystems), and quantification of gene expression was performed using a standard curve and normalized to *Actb* expression levels. qRT-PCR primer sequences can be found in **Table 1**.

**Table 1:**
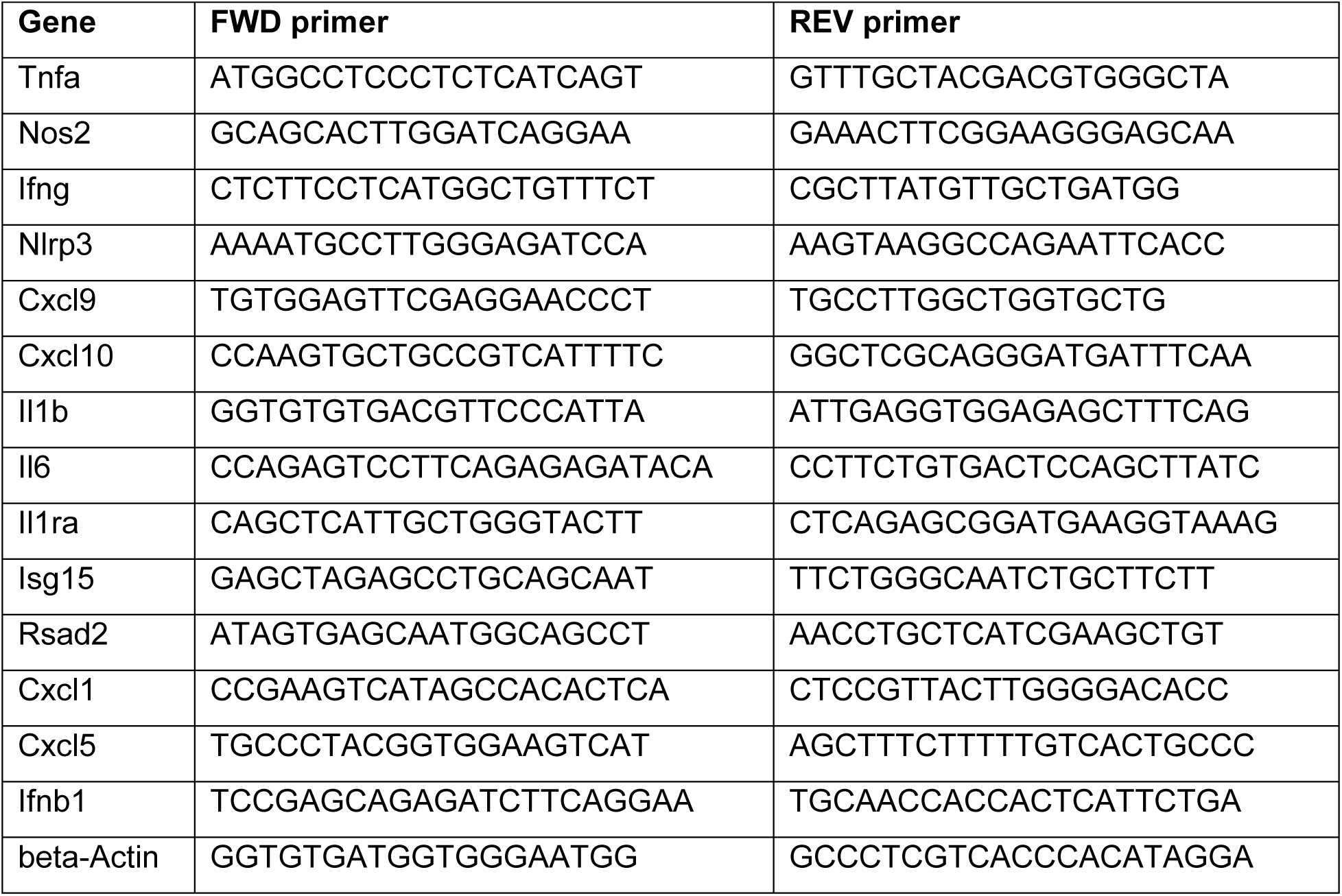
qRT-PCR primer sequences.

### Flow cytometry

For *ex vivo* analysis of lung cell populations at 21 days post Mtb infection (WT (n=6) and Polg^D257A^ (n=8)), the right inferior lobe was isolated and washed twice in 1X PBS. The lung was then minced and digested in digestion buffer (70 μg/ml Liberase (Roche) and 50 μg/ml DNASE I (manufacturer) in RPMI 1640 (HyClone)) for 30 min at 37°C 5% CO_2_. Single suspensions were achieved by consecutively filtering homogenates through 70 μM and 40μM cell strainers. Live dead staining was performed using Ghost dye 510 (Tonbo). Fc receptors were blocked using CD16/CD32 monoclonal antibody (eBiosciences). Cells were stained with antibodies against surface proteins CD11b (BV421, BDBiosciences), CD11c (BV605, BioLegend), CD45 (BV785, BioLegend), CD170 (eFluor-488, eBiosciences), CD43 (PE, BD Biosciences), Ly6G (PerCP-Cy5.5, eBiosciences), Ly6C (APC, eBiosciences), CD206 (APCeFLuor-700 eBiosciences), CD86 (APCeFluor-780, eBiosciences). Cells were washed 2 times before fixing in 4% PFA for 15 min at RT. Following fixation cells were washed twice with 1X PBS and incubated overnight at 4°C. Total lung cell counts were based on live single cells (Ghost low/-) in 200 mL or 1/3 lung lobe. FSC/SSC was used to differentiate macrophage (CD45^+^ CD11b^+^ Ly6G^-^ CD11c^low/-^ SSC^hi^ FSC^mid^) and monocyte ((CD45^+^ CD11b^+^ Ly6G^-^ CD11c^low/-^ SSC^lo^ FSC^mid^) Flow cytometry was performed on the LSR Fortessa X20, and FCS Express software was used for post-acquisition analysis. See Figure S1A for flow gating strategy.

### Protein quantification by immunoblot

Cells were washed with 1XPBS and lysed in 1X RIPA buffer with protease and phosphatase inhibitors (Thermo Scientific), with the addition of 1 U/mL Benzonase (Millipore) to degrade genomic DNA. Proteins were separated by SDS-PAGE Any kD mini-PROTEAN TGX precast gels (Bio-Rad) and transferred to 0.45μm nitrocellulose membranes (GE Healthcare). Membranes were blocked for 1 hr at RT in TBS with 5% BSA. Blots were incubated overnight at 4°C with the following antibodies: β-TUBULIN (Abcam, ab179513) and VIPERIN (Millipore, MABF106). Membranes were incubated with appropriate secondary antibodies for 2 hrs at RT prior to imaging on a LiCOR Odyssey Fc Dual-Mode Imaging System. For protein isolated from tissue, lung homogenates were lysed in 1% SDS lysis buffer and then boiled for 30 min at 95 °C. Protein per sample was quantified using a bicinchoninic acid assay (BCA) (Thermo Fisher Scientific, PI23235). 30 μg of total protein was loaded into 10% SDS-polyacrylamide gels and transferred onto 0.22 μM polyvinylidene difluoride membranes, dried, and then incubated overnight at 4 ᵒC rolling with the following primary antibodies in 1x PBS containing 1% casein: VIPERIN (Abcam, ab107359), IL-1β (Cell Signaling Technology, 12242), NLRP3 (Cell Signaling Technology, 15101), GAPDH (ProteinTech, 60004-1-Ig), and ACTIN (ProteinTech, 66009-1-Ig). Next, membranes were washed and incubated with horseradish peroxidase (HRP)-conjugated secondary antibodies for 1 hour at room temperature. Luminata Crescendo Western HRP Substrate (Millipore, WBWR0500) was used to visualize proteins.

### Histopathology

Mtb infected mouse lungs were fixed with 10% neutral formalin and processed, embedded in paraffin, and cut into 5 μM sections and stained with hematoxylin and eosin (H&E) or acid-fast stain (AFB) (AML Laboratories). A boarded veterinary pathologist performed a blinded evaluation of lung sections for inflammation. To quantify the percentage of lung fields occupied by histiocytic inflammatory infiltrates, scanned images of a lobe of each lung were analyzed using QuPath Bioimage analysis v 0.4.3 to determine the total cross-sectional area of inflammatory foci per total lung cross sectional area. Additional criteria were evaluated by dividing the digital images into 500 × 500 μm grids and counting the percentage of squares containing neutrophils and neutrophils arranged in clusters (> 5 neutrophils in close approximation). Necrotic debris represents the nuclear remnants of dead cells. Necrotic debris was classified as extracellular, circular, densely basophilic structures which were variably sized, though always smaller than an erythrocyte (less than 6 μm), and frequently seen in association with neutrophil clusters. WT and Polg^D257A^ mice lung sections from were divided into 500 × 500 μm grids and evaluated by counting the percentage of squares based on the presence or absence of necrotic debris.

### Quantification of acid-fast bacteria (AFB) staining

To directly quantify the bacillary burden within regions of inflammatory infiltrates, a certified veterinary pathologist used QuPath Bioimage analysis v 0.4.3 to annotate the total area of inflammatory regions in each acid-fast stained lung section. Subsequently, a script was used to randomly sample ten 100 x 100 μm square regions of interest within these areas. In these regions, a reviewer blinded to the study details manually counted the carbolfuchsin-positive bacilli.

### Statistical analysis

All data are representative of two or more independent experiments with n=3 or greater unless specifically noted in the figure legends. For all quantifications, n represents the number of biological replicates, either number of wells containing cells or number of mice. Error bars represent SEM. For *in vitro* assays, statistical significance was determined using either a two-tailed Student’s unpaired T test or two-way ANOVA with Tukey’s post hoc test. For *in vivo* mouse infections, significance was determined using a Mann-Whitney U test based on the assumption that samples (mice) followed a nonnormal distribution. The specific statistical test used to determine significance for each experiment is listed at the end of the figure legends. All calculations of significance were determined using GraphPad Prism Version 10 Software expressed as P values. The threshold for significance was determined by a p value of < 0.05, and annotated as * = p<0.05, ** = p<0.01, *** = p<0.001, **** = p<0.0001. For *in vivo* infection mouse experiments, we estimated that detecting a significant effect requires two samples to differ in CFUs by 0.7e^10. Using a standard deviation of 0.3e^10 for each population, we calculated that a minimum size of 5 age- and sex-matched mice per group per time point is necessary to detect a statistically significant difference by a t-test with alpha (2-sided) set at 0.05 and a power of 80%. Therefore, we used a minimum of 5 mice per genotype per time point to assess infection-related readouts.

