## Supplemental Files for "Necrosis drives susceptibility to *Mycobacterium tuberculosis* in Polg^D257A^ mutator mice"

A

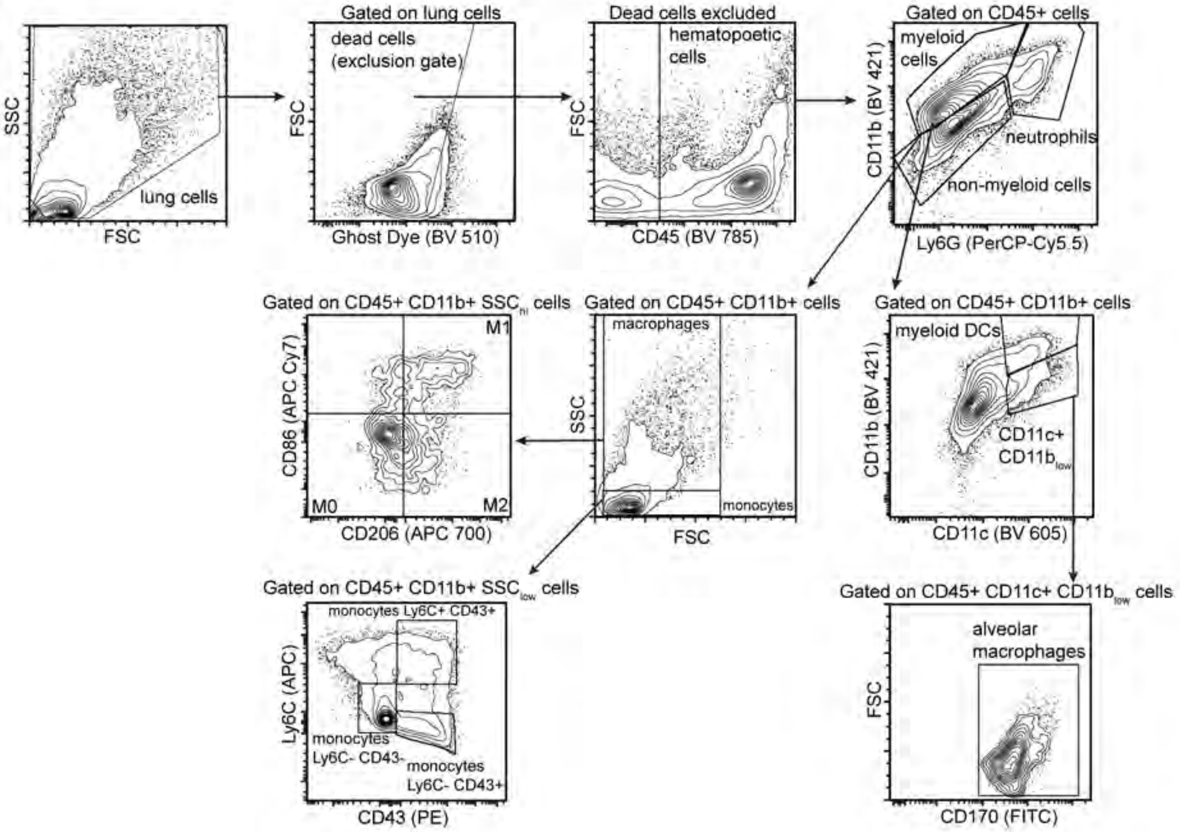
lung cells
SSC
FSC
Gated on lung cells
dead cells
(exclusion gate)
FSC
Ghost Dye (BV 510)
Dead cells excluded
hematopoetic
cells
FSC
CD45 (BV 785)
Gated on CD45+ cells
myeloid
cells
neutrophils
non-myeloid cells
CD11b (BV 421)
Ly6G (PerCP-Cy5.5)
Gated on CD45+ CD11b+ SSChi cells
M1
M0
M2
CD86 (APC Cy7)
CD206 (APC 700)
Gated on CD45+ CD11b+ cells
macrophages
monocytes
SSC
FSC
Gated on CD45+ CD11b+ cells
myeloid DCs
CD11c+
CD11blow
CD11b (BV 421)
CD11c (BV 605)
Gated on CD45+ CD11b+ SSClow cells
monocytes Ly6C+ CD43+
monocytes
Ly6C- CD43-
monocytes
Ly6C- CD43+
Ly6C (APC)
CD43 (PE)
Gated on CD45+ CD11c+ CD11blow cells
alveolar
macrophages
FSC
CD170 (FITC)

Figure S1

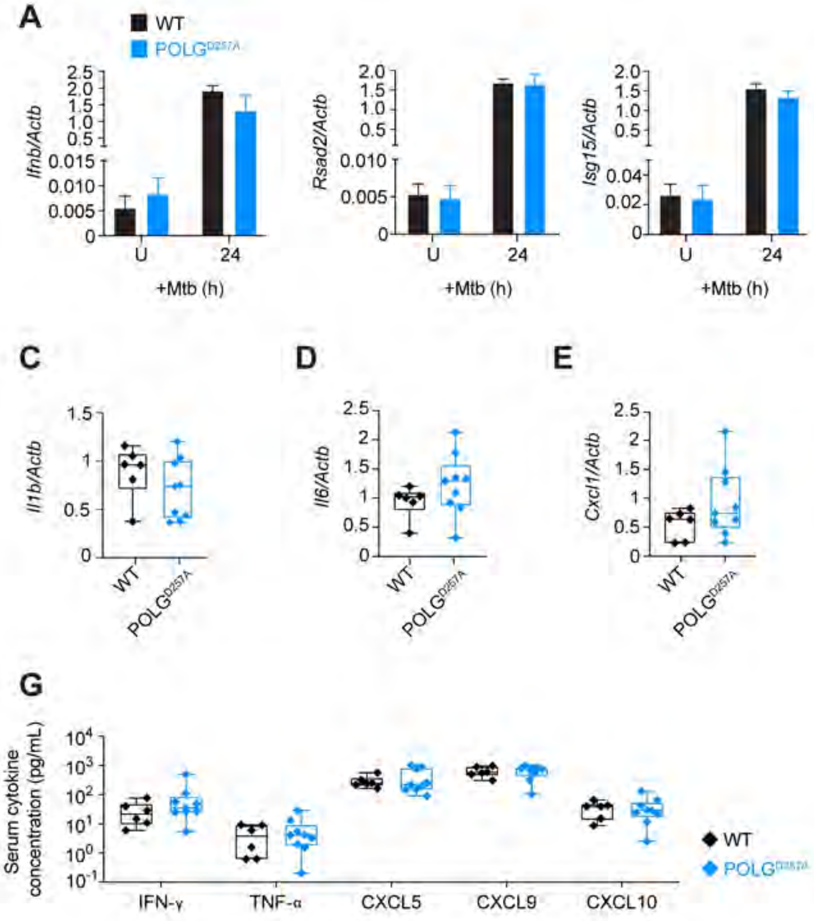
A
WT
POLG D257A
Ifnb/Actb
Rsad2/Actb
Isg15/Actb
U
24
+Mtb (h)
B
+Mtb
U
24h
VIPERIN
TUBULIN
VIPERIN/TUBULIN
C
Il1b/Actb
D
Il6/Actb
E
Cxcl1/Actb
F
Cxcl5/Actb
G
Serum cytokine concentration (pg/mL)
IFN-γ
TNF-α
CXCL5
CXCL9
CXCL10

Figure S2

**Figure S1. Lung immune cell analysis of WT and Polg$^{D257A}$ mutator mice during Mtb infection.**

**(A)** Flow gating strategy for analysis of lung innate immune cell populations during Mtb infection *in vivo*. FCS Express software was used for post-acquisition analysis.

**Figure S2. Polg$^{D257A}$ mutator mice do not show differences in interferon stimulated genes (ISGs) during Mtb infection**

**(A)** *Ifnb*, *Rsad2*, and *Isg15* transcripts analyzed by qRT-PCR in untreated and 24h Mtb infected (MOI=5) WT and Polg$^{D257A}$ macrophages. Transcripts were normalized to *Actb* transcript levels. **(B)** Immunoblot analysis of VIPERIN protein levels in untreated and 24h Mtb infected (MOI=5) WT and Polg$^{D257A}$ macrophages. Protein levels normalized to loading control $\beta$-TUBULIN. Lung *Il1b* **(C)**, *Il6* **(D)**, *Cxcl1* **(E)**, *Cxcl5* **(F)** transcripts in WT and Polg$^{D257A}$ mice analyzed by qRT-PCR at day 21 post- Mtb infection. Transcripts were normalized to *Actb* transcript levels. **(G)** Circulating serum cytokines in WT and Polg$^{D257A}$ mice at day 21 post- Mtb infection.

Statistical analysis: *$p < 0.05$, **$p < 0.01$, ***$p < 0.001$, ****$p < 0.0001$. Statistical significance determined by using (A,B) two-tailed Student's t test and (C-G) Mann-Whitney U test. Error bars represent SEM.
